# Leveraging epigenetic signatures to determine the cell-type of origin from long read sequencing data

**DOI:** 10.1101/2024.06.03.597114

**Authors:** Eilis Hannon, Jonathan Mill

**Affiliations:** Department of Clinical and Biomedical Sciences, University of Exeter Medical School, University of Exeter, Barrack Road, Exeter, Devon, EX2 5DW, UK

**Author notes:** **Corresponding author: Eilis Hannon, Department of Clinical and Biomedical Sciences, University of Exeter Medical School, RILD building, Royal Devon & Exeter Hospital, Barrack Road, Exeter. EX2 5DW. UK.**.

## Abstract

DNA methylation differs across tissue- and cell-types with important implications for the analysis of disease-associated differences in tissues such as blood. To uncover the biological processes affected by epigenetic dysregulation, it is essential for epigenetic studies to generate data from the appropriate cell-types. Here we propose a framework to do this computationally from long-read sequencing data, bypassing the need to isolate subtypes of cells experimentally. Using reference data for six common blood cell-types, we evaluate the potential of this approach for attributing reads to specific cells using sequencing data generated from whole blood. Our analyses show that cell-type can be accurately classified using small regions of the genome comparable in size to those generated by long-read sequencing platforms, although the accuracy of classification varies across different regions of the genome and between cell-types. We found that for approximately one third of the genome it is possible to accurately discriminate reads originating from lymphocytes and myeloid cells with the prediction of more specialised subtypes of blood cell-types also encouraging. Our approach provides an alternative computational method for generating cell-specific DNA methylation profiles for epigenetic epidemiology, accelerating our ability to reveal critical insights of the role of the epigenome in health and disease.

## Introduction

An important advance in our understanding of the genetic risk factors for complex traits such as diabetes, Alzheimer’s disease and cardiovascular disease, is the conclusion that most variants primarily exert their effects by influencing gene expression. This has focused attention on understanding the role of the epigenome, which encompasses a diverse number of chemical modifications to DNA and nucleosomal histone proteins that directly influence gene expression, in the aetiology of these traits (Li et al. 2019; Battram et al. 2022). The underlying properties of the epigenome make it particularly relevant to human disease; it is highly dynamic, varying across development, between cell-types and in response to the environment. However, this fundamental difference compared to the relative stability of genetic variation, means that careful consideration to study design is required when identifying epigenetic variation associated with complex traits (Relton and Davey Smith 2010; Mill and Heijmans 2013).

The most commonly studied epigenetic modification in human health and disease is DNA methylation (Murphy and Mill 2014; Campagna et al. 2021), which involves the addition of a methyl group to the fifth carbon position of cytosine. While most studies have used microarray technologies to profile DNA methylation (DNAm) across the genome, it is the first modification that can be reliably characterised with next generation sequencing technologies in large numbers of samples (Simpson et al. 2017). Because the epigenome orchestrates the gene expression changes underpinning cellular differentiation during development, the DNAm profile of a sample is primarily defined by the tissue or cell-type it originates from (Hannon et al. 2015; Roadmap Epigenomics Consortium et al. 2015; Chen et al. 2016; Hannon et al. 2021; Salas et al. 2022). The choice of tissue for profiling, therefore, has a profound effect on not only the exact results obtained, but additionally, the nature of conclusions that can be drawn from the analysis of differential DNAm. Profiling the primary affected tissue, e.g. the cortex for Alzheimer’s disease, is pertinent for making inferences about how DNAm variation contributes to the aetiology of a particular trait.

A major caveat with profiling modifications to DNA isolated from ‘bulk’ tissue, (e.g. whole blood or a dissected section of the cortex) is that it represents a heterogeneous mix of different cell-types. Each of these cell-types has a specific DNAm profile, with the resulting profile of the bulk tissue being an aggregate of those from the constituent cell-types. As the proportion of individual cell-types within a sample can vary across individuals, systematic differences in cellular proportions that correlate with the phenotype of interest may manifest as differences in the overall DNAm profile (Jaffe and Irizarry 2014). Knowledge of the composition of cell-types within each sample is critical for minimising false positive associations in epigeneome-wide association studies (EWAS); to counter this analyses often include quantitative covariates that capture the cellular composition of each sample (Jaffe and Irizarry 2014). While this may prevent false positive associations driven by cell-type differences between groups, it does not enable us to identify the cell-specific changes that might be important in disease. In addition, subtle changes or differences in rarer cell-types may go undetected as they compete against the background signal from more abundant cell-types. Knowing which cell-type is the context for differences in DNAm (and potentially by proxy gene expression) is critical for determining the genes and biological processes associated with specific complex traits and ultimately developing novel targets for preventing and treating disease. It is not just epigenetics where the cellular context is important; pathogenic mutations associated with cancer occur in specific cell lineages and identifying these can be valuable for diagnosis.

The case for cell-specific DNAm profiling is compelling, however generating these data is challenging. It is expected that entirely new experimental datasets will be needed, generated from purified cell populations isolated using physical isolation methods such as fluorescent-activated cell sorting (FACS), a challenging, costly and timely endeavour. While we and others have started to generate reference data for specific cell populations, sample sizes remain small and analyses underpowered (Chen et al. 2016; Gasparoni et al. 2018; Shireby et al. 2022). As an alternative, a number of computational solutions have been proposed bypassing the need to isolate cells prior to DNAm profiling. These methods use the estimated cellular composition profile in tandem with bulk DNAm profiles to either mathematically derive cell-specific profiles prior to analysis (Rahmani et al. 2019) or implement a more complex regression model, with interaction terms, to determine the cell-specificity of differences associated with the outcome (Zheng et al. 2018). Both approaches are inherently dependent upon knowing or estimating the cellular composition of the samples. The accuracy of the quantitative variables that capture the abundance of each cell-type, which could be inferred either experimentally or computationally, influences both the power and accuracy of downstream association analyses. Furthermore, these methods are applied to each genomic position separately; the complexity of the disaggregation calculation will depend on how cell-specific the profiles are at that position. For example, at a position where methylation is exclusive to one cell-type, any difference in DNAm measured in bulk tissue can be attributed to that cell-type. However, where multiple cell-types share a methylation status at a specific site, it becomes more challenging or perhaps even impossible to assign the signal to a particular cell-type. Minimal experimental evidence has been provided thus far to assess what proportion of the genome these approaches can be used to accurately determine cell-specific DNAm differences associated with complex traits. Quantifying the specificity of DNAm variation across blood cell-types, we previously demonstrated that only in the minority of cases was a single cell-type responsible for the variation observed in whole blood (Hannon et al. 2021). While similar analyses in a wide range of different bulk tissues will be needed to generalise this finding, it suggests that for the majority of the genome, the way cell-specific DNAm profiles combine into a bulk tissue profile is not simple and therefore these deconvolution methodologies will be associated with high error rates.

In this manuscript, we propose an innovative alternative computational approach that has been enabled by the availability of long read sequencing technologies and is not contingent on knowing the cellular composition of a sample. The theory behind our approach centres on the fact that each individual sequencing read originates from a single cell. There are two, equally critical technological advances that we exploit in this methodology: i) the generation of long reads capturing thousands of bases of sequence and ii) the ability to directly detect DNAm at base-pair resolution in parallel to the genetic sequence (Lucas and Novoa 2023). Harnessing these two advances together means that within each long-read a significant proportion of the DNAm profile of that cell is captured. The principle of our approach is to use this information to classify the cell-type each individual read originates from. After classification we can then group reads by cell-type prior to aggregating into cell-specific epigenetic profiles. In this way, we can generate cell-specific DNAm datasets from a bulk tissue long read sequencing experiment for downstream EWAS analyses and integrate data on phased genetic variation and DNA modifications from the same molecule.

The primary objective of this study is to determine the feasibility of this approach, which theoretically holds great promise, but may be limited by two methodological caveats. First, existing knowledge of cell-type-specific DNAm profiles leverages information across the genome by incorporating multiple regions to characterise the signature of each cell-type. As each individual read may contain only a few thousand kilobases, it is unknown if the frequency of cell-specific signatures across the genome is sufficient to ensure that a single long read would contain enough discriminating information. Second, most DNAm profiling methods are quantitative, as they quantify methylation status across a population of cells. For a specific genomic position, a continuous value is obtained by aggregating across multiple cells and can be used to make relative comparisons between conditions. With a continuous measure, depending on the magnitude of variation within a cell-type, there is the potential to distinguish an unlimited number of different cell-types from a single genomic position. However, for each individual DNA molecule from an individual cell the methylation status is binary. This loss of information makes the classification problem more challenging, and means more genomic positions will be required in any classifier to compensate. From a single genomic position, it is only possible to classify at most two groups of cell-types. To classify three or more cell-types will require multiple genomic positions and therefore is not possible with short read sequencing.

In this paper, we leverage existing reference epigenetic data, specifically genome-wide maps of DNA methylation (DNAm) in purified blood cell-types (Hannon et al. 2021; Loyfer et al. 2023), in tandem with supervised machine learning classification algorithms to determine the feasibility of computationally assigning long sequencing reads to different cell-types. We have focused this investigation on classifying the major blood cell-types as this represents the predominant tissue profiled in epigenetic epidemiology. We are particularly interested in addressing three key questions that underpin the feasibility of our proposed approach. First, can we classify cell-type based on epigenetic profiles spanning just a few thousand bases of sequence? Second, is the distribution of cell-specific signatures through the genome frequent enough that each long read will contain at least one informative position? Third, is there enough information within a long read to classify cell-types when measuring a binary modification such as DNAm?

## Results

### Cell-type can be accurately classified using segments of less than 20kb of the genome

It is well established that genome-wide DNAm profiles can be used to define cell-type identity (Hannon et al. 2021). However, it is unknown whether similar discrimination can be achieved with a smaller snapshot of the epigenome. Our first objective was to determine whether it was feasible to predict cell-type using a subset of the genome (e.g. a 20kb segment from a single chromosome) and investigate how small this region could be. To this end we used DNAm data for five purified blood cell-types generated using the Illumina EPIC microarray (Hannon et al. 2021) (**Supplementary Table 1**). This technology only captures ∼3% of the CpGs in the human genome, which is the sequence context where DNAm is concentrated. Given the sparse coverage in our training dataset, our analyses likely overestimate the minimum window required. From 783,501 autosomal DNAm sites, we tested all possible classifiers consisting of at least 5 DNAm sites and up to 20kb of the human genome, a total of 3,427,657 distinct classifiers. Here we use the term classifier to refer to a unique combination of contiguous DNAm sites, which are the features of the algorithm. Classifiers had a median of 11 (IQR = 7 - 18) sites, spanned a median 9,779bp (IQR = 4,467 – 14,901bp) of the genome, with a median of 1 DNAm site per 663bp (IQR = 328 – 1,166bp)(**Supplementary Figure 1**). Each classifier was trained using four different machine learning algorithms, support vector machine (SVM), Naïve Bayes, K nearest neighbours (KNN), and Random Forest.

Generally, classifiers were able to predict cell-type accurately (**Figure 1**). There was little to choose between the performance of the Random Forest (mean accuracy = 0.80, SD = 0.14), the Naïve Bayes (mean accuracy = 0.79, SD = 0.14), and the KNN (mean accuracy = 0.78, SD = 0.15) algorithms, although the SVM algorithm was associated with a markedly reduced accuracy score (mean accuracy = 0.70, SD = 0.15)(**Figure 1A**). While it is encouraging that virtually all classifiers outperform random assignment of labels, which would be associated with a mean accuracy of 0.2, for our purposes of generating cell-specific epigenetic profiles, we require a higher level of accuracy. At a minimum threshold of 90% samples correctly assigned (i.e. accuracy > 0.9), there were 1,070,740 classifiers (31%) that surpassed this threshold with at least one algorithm (**Figure 1B**). While Random Forest was associated with the highest number of accurate classifiers (996,831, 29%) there were 73,909 (2%) classifiers that passed this threshold only with an algorithm other than Random Forest, highlighting in a subset of cases the benefit of implementing multiple machine learning algorithms. Typically for a classifier, if one algorithm produced an accurate prediction for a region of the genome, all the algorithms did well, suggesting that the selection of DNAm sites is a more important factor than algorithm (**Supplementary Figure 2**). For the purposes of simplifying the presentation of the results, and to retain as many classifiers as possible, for each classifier we took the single best performing algorithm; this approach marginally increased the mean accuracy across all classifiers to 0.81 (SD = 0.14).

**Figure 1.**
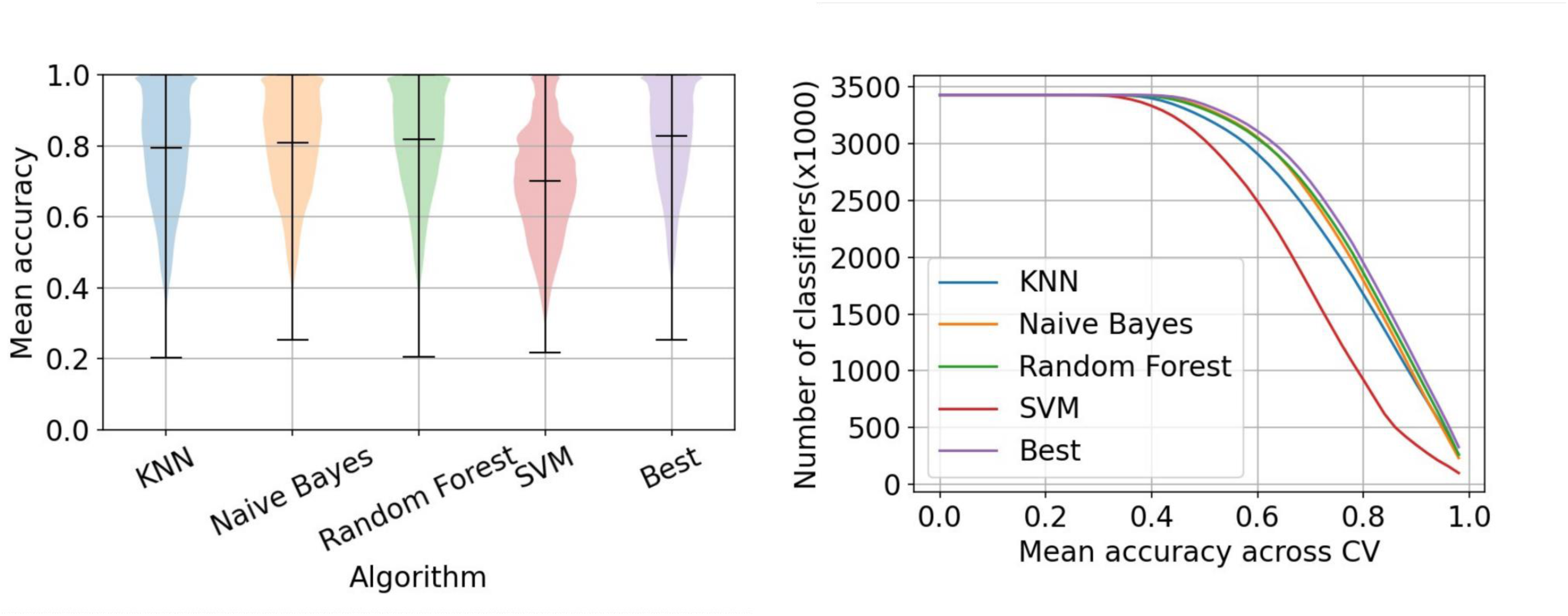
Cell-type can be accurately classified from segments of the genome less than 20kb. A) A Violin plot of the distribution of accuracy scores across classifiers, grouped by machine learning algorithm. B) Line graph of number of classifiers (y-axis) with a minimum mean accuracy (x-axis), where each line represents a different machine learning algorithm.

To understand the variation in accuracy across classifiers, we grouped them based on the properties of the DNAm sites they contained and summarised the accuracy statistics across these groups (**Figure 2** and **Supplementary Figure 3**). We observed that increasing the number of sites in the classifier does lead to improved accuracy, although this relationship is not linear. There is a dramatic improvement when increasing from 5 to 20 CpGs, with more subtle improvements up to 50 CpGs, there is then a more dramatic improvement per CpG up to 80 sites after which the relationship plateaus (**Figure 2A**). It is also noticeable that with more sites in the model, there is less variability in the performance of the classifiers and the minimum performance level can be considered as very accurate. We also observe a non-linear monotonic relationship between accuracy and the size of the genomic segment captured by the classifier (**Figure 2B**). Again, there is an initial dramatic improvement as segments increase in size up to about 2.5kb, after which there are decreasing gains as the graph starts to plateau. Of note, at 20kb - the maximum window we considered - the graph has yet to reach a plateau, and the variability across classifiers is still large. Together, this suggests that it is the number of features in a genomic segment rather than size of the segment that is more important. Finally, we looked at the effect of genomic density of sites in the segment (**Figure 2C**) finding a non-linear relationship, whereby after an initial dramatic improvement, maximum average performance was obtained when sites were located on average every 650-700bp, and performance decreased as the mean gap between sites increases. This suggests that cell-specific signatures cluster around CpG dense genomic regions, supporting previous data demonstrating that cell- and tissue-specific signatures do occur at CpG islands(Illingworth et al. 2008). These results may also indicate that having multiple CpGs close together (that presumably have a correlated methylation profile), improves the sensitivity of detecting differences by increasing the accuracy of the estimate of DNAm level in that genomic region.

**Figure 2.**
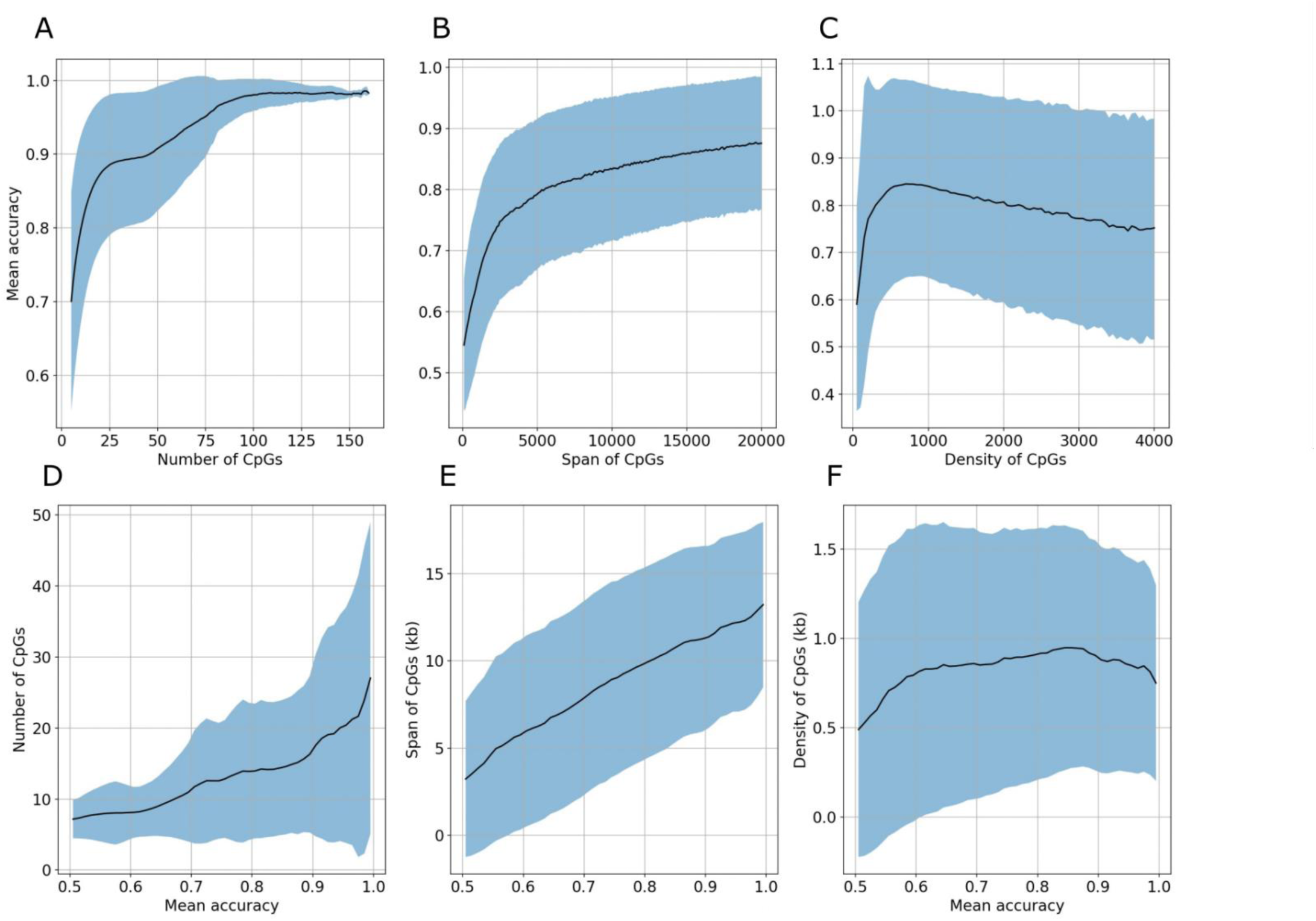
Classifier accuracy improves with more CpGs closer together. The top row (**A-C**) presents summaries of the accuracy for classifiers grouped by CpG properties, while the bottom row (**D-F**) presents summaries of CpG propoerties for classifiers grouped by accuracy. Where the CpG properties consdiered where **A,D**) number of CpGs, **B,E**) genome range of CpGs and **C,F**) the mean gap between CpGs. Each panel shows the mean as a (solid) line with +/- 1 standard deviation shown by the shaded area. In all panels the grouping variable is on the x-axis and the summary statistic on the y-axis. Accuracy is defined at the proportion of samples predicted correctly, summarised for each classifier as the mean accuracy score across 15 cross-fold validation iterations, for each classifier we took the best accuracy metric across the four algorithms.

While it is logical that increasing the number of sites included in a classifier would lead to improvements in accuracy, as the model has more information (i.e. features) to use for discrimination, it does not necessarily follow that the most accurate classifiers require lots of DNAm sites. To determine how much information an accurate classifier needs, we instead grouped classifiers based on their accuracy metrics and summarised the metrics relating to the number of sites they contained (**Figure 2** and **Supplementary Figure 4**). To obtain an accuracy of at least 0.9, a mean of 17 sites (SD = 12 CpGs)) spanning 11.4kb (SD = 5.2kb) at a density of 1 site per 880bp (SD = 634bp) is needed. Together, this demonstrates that where cell-specific DNAm signatures exist, not many sites are required for an accurate classifier, but by increasing the size of the genomic segment the likelihood that the classifier incorporates one of these signatures is increased.

### Accurate classifiers cluster by genomic position with large gaps in between

Given the sliding window methodology we used to test all possible combinations of sites, many of the classifiers are nested within each other. It is logical that all classifiers from the same genomic region with overlapping feature sets will have correlated accuracy statistics. In order to quantify what proportion of the genome contains an accurate classifier, we collapsed predictive classifiers into a set of non-overlapping regions, where one region likely contains multiple classifiers (**Supplementary Figure 5**). At an accuracy threshold of 0.9, the 1,070,740 classifiers reduced to 8,264 genomic regions. Each region contained a mean of 130 classifiers (SD = 624) and spanned a mean of 38.8kb (SD = 32.4kb) of the genome. In total they represented 11% of the autosomal human genome with the mean gap between regions was 309 kb (SD = 1.1mb). This indicates that while there is promise in our strategy of predicting cell-type from a single long read, for reads shorter than 20kb it is only viable for a fraction of the genome.

### Simplifying the prediction problem to classifying a single cell-type improves performance

While one strategy for increasing the proportion of the genome that contains an accurate cell-type classifier is to increase the size of the genomic segment, an alternative approach is to simplify the classification problem by predicting fewer cell-types. We tested the potential gains of this hypothesis, by re-training classifiers to predict a maximum of two cell-types. We did this in two ways. First, grouping blood cell-types into more general, higher order categories (i.e. lymphocyte or not). Second, by training a series of binary predictors for each cell-type (i.e. granulocyte or not). This dramatically improved the performance of the predictors with mean accuracies across all classifiers ranging from 0.91 – 0.98 (**Figure 3**).

**Figure 3.**
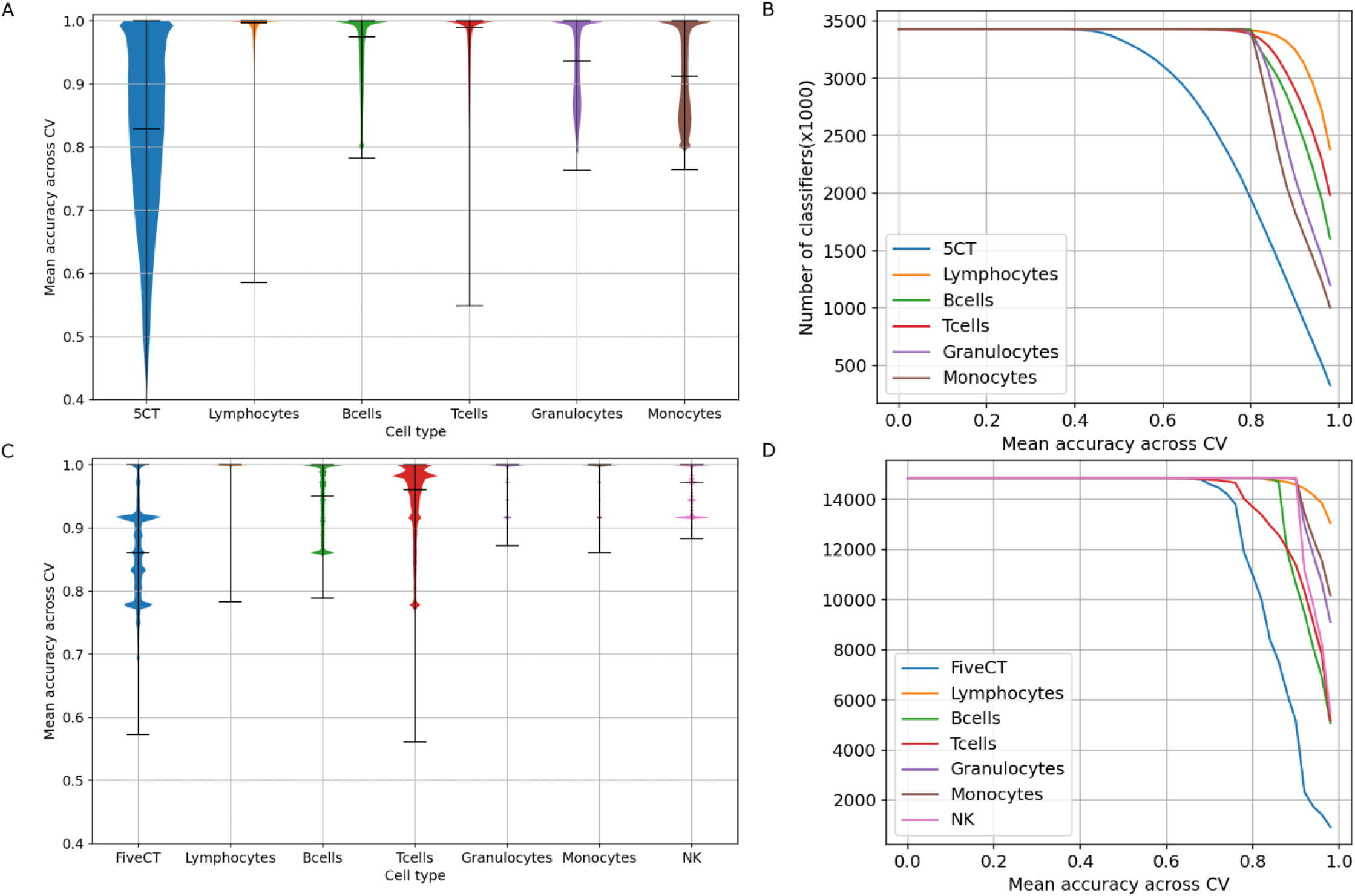
Binary cell-type classifiers outperform classifiers predicting multiple blood cell-types simultaneously. Plotted are a summary of the accuracy statistics for the simplified binary classification problem of one cell-type vs all others. Accuracy is defined at the proportion of samples predicted correctly, summarised for each classifier as the mean accuracy score across 15 cross-fold validation iterations, for each classifier we took the best accuracy metric across the four algorithms. Panels A & B were trained used the EPIC microarray reference cell-type data, while panels C & D were trained using the WGBS reference cell-type data. Panels A & C show violin plots of the distribution of accuracy scores across binary classifiers. Panels B & D show lines graph of the number of classifiers (y-axis) with a minimum mean accuracy (x-axis), where each line represents a different classifier for a different cell-type.

With the classifiers distinguishing higher-level cell-type groupings of lymphocytes vs myeloid cells more effectively (mean accuracy = 0.98, SD = 0.04) than predicting more specialised blood cell-types such as granulocytes or not. As with the more complex classification problem, we observed that the more data in the model (e.g. number of sites or length of genomic window) that was considered, the accuracy of the prediction increased.

Furthermore, for these binary classification problems, the performance plateaued sooner (i.e. with few sites) than when trying to predict five cell-types simultaneously, and plateaued at higher mean accuracy scores (∼> 0.95) (**Supplementary Figure 6**). For example, focusing on only the classifier trained to distinguish lymphocytes from other blood cell-types, a mean of just 7 CpGs or 4.7kb of sequence are needed to obtain an accuracy of 0.9, less than half of the requirements for the five-cell-type prediction problem (**Supplementary Figure 7**). These properties vary across the binary classifiers for each cell-type, with the monocytes being the most challenging to predict and requiring a mean of 14 sites or a genomic segment of 10.3kb to get a mean accuracy of > 0.9, only slightly less than the mean requirements of the classifiers trained to predict five cell-types simultaneously.

As with the five cell-type classifiers, we collapsed the binary classifiers into non-overlapping regions to quantify the proportion of the genome that contained an accurate classifier. For the lymphocyte vs myeloid classifiers, at an accuracy threshold of 0.9, 3,249,569 classifiers were reduced to 16,170 regions. As well as there being approximately twice as many regions as obtained when classifying five cell-types simultaneously, the regions were on average longer with a mean size of 46.9 kb. Therefore in total they cover more than a two-fold increase in proportion of the human genome (0.11 vs 0.26). Considering the binary classifiers for the other five specialised blood cell-types, we observed slightly fewer genomic regions (between 12,736 and 15,137), with a similar mean size (42.2 – 46.8kb) which covered a larger proportion of the human genome (0.18 – 0.24) than the classifiers for five cell-types (**Supplementary Figure 8**). Together, this demonstrates that there are significant gains from reframing the objective to train binary cell-type predictors and suggests that highly predictive signatures for individual cell-types are small enough and frequent enough across the human genome to be located within a single long read.

### Cell-type classifiers are associated with higher accuracy when trained using the denser coverage of CpG sites available from whole genome bisulfite sequencing

A second strategy for increasing the coverage of accurate cell-type classifiers is to increase the resolution of sites in the training data with a technology that covers a larger proportion of sites in the genome, such as whole genome bisulfite sequencing (WGBS). The trade-off here is typically smaller sample sizes and lower sensitivity in the estimation of DNAm level due to read depth(Seiler Vellame et al. 2021). Taking advantage of a publicly available DNA methylation atlas of human cell-types (Loyfer et al. 2023) (**Supplementary Table 1**), we repeated our analyses with this WGBS data as training data. This increased the number of classifiers to 14,849,740 consisting of a median of 18 sites (IQR = 10 - 31) covering a median of 9.4 kb (IQR = 4.4 – 14.6 kb) with a median of 1 site per 397bp (IQR = 211 – 714) (**Supplementary Figure 9**). Applying these to the two-cell-type binary classification problems we found a mean accuracy of between 0.94 – 0.99 (**Figure 3C**), marginally higher than when using the EPIC array derived profiles as training data. However, the percentage of models with accuracy > 0.9 was higher. More than 98% of the classifiers predicting either lymphocyte, NK, granulocyte or monocytes had an accuracy > 0.9 with only 72% for B-cells and 77% for T-cells respectively. While there was variation in the performance of the binary classifiers for the different cell-types, the rank order of accuracy statistics differed depending on whether the classifiers were trained with the EPIC array data or WGBS data. For example, monocytes were associated with the worst performance when trained with the EPIC array data but the best performance when trained with the WGBS data. This suggests factors specific to the generation of each reference profile, e.g. purity, data quality or omission of informative sites for that cell-type, are introducing variation in performance and we cannot necessarily generalise about which blood cell-types are easier or harder to classify. Rather than the binary classification problem, getting the classifiers to try the more complex problem of predicting all five blood cell-types present in the WGBS training dataset simultaneously reduced the mean accuracy to 0.86 (SD = 0.07).

There was an interesting non-linear relationship between the number of sites and accuracy of WGBS trained classifiers, whereby after the anticipated gain in performance as number of CpGs increases, there was a drop in mean performance for classifiers with between 50 and 220 CpGs (**Supplementary Figure 10**). This was mirrored across the classifiers for all cell-types. This might indicate that there are nuances in the distribution of sites across the genome, or that as the number of sites increases the probability of one of those sites having an inaccurate estimate of DNAm level - due to either low read depth or stochasticity in the sequencing process - also increases. Considering performance against the size of the genomic segment represented by the classifier, there were dramatic improvements up to 1kb, with more moderate increments for classifiers longer than this. Again, performance improved sharply to a peak, before tailoring off more gradually as a function of site density with peak accuracy obtained at a density of 1 site per 500bp. Of note, when classifiers are grouped by accuracy statistics (**Supplementary Figure 11**), binary classifiers with accuracy > 0.9 require a mean of 7-21 sites, slightly higher on average than classifiers trained with the EPIC array data; this difference is probably attributable to lower sensitivity in estimating in DNAm level in sequencing data. However, these sites are contained within smaller genomic segments of ∼4-8.5kb reflecting the fact that the WGBS data has a denser coverage of sites. Ultimately, this indicates that it is beneficial to train with genuine genome-wide DNAm profiles if the aim is to constrain the genomic segment used to predict cell-type. In line with the results from the classifiers trained with the EPIC array reference data, more sites over a bigger genomic window were required for similar levels of performance when predicting all five cell-types simultaneously compared to the binary classification problem (mean 25 sites, 10.6 kb for accuracy > 0.9).

Collapsing these results into non-overlapping regions, despite many more classifiers with accuracy > 0.9, the total number of regions was similar to that when training with the EPIC array data. For example, for lymphocytes, 14,585,430 predictive classifiers (accuracy > 0.9) collapsed into 14,927 regions, however the regions are on average longer with a mean size of 73.1kb. Altogether, therefore, they capture an increased proportion of the genome, 0.38 (**Supplementary Figure 12**), demonstrating that the more comprehensive profiling of CpG sites across the genome yields a higher fraction of usable data when classifying long read data into distinct cell-types.

### Larger genomic regions are required to predict cell-type from individual read level binary DNA methylation status data

Having demonstrated the potential of using a subset of the genome to predict cell-type, we next wanted to assess how this approach would work when applied to long read sequencing data. One critical difference is that at the single read level, DNAm status is binary as it originates from a single cell, whereas in the classifier models we have trained thus far, DNAm status is quantified as a proportion (i.e. a continuous value), representing the mean DNAm status across a population of cells. This shift in the input data from continuous to binary will reduce the amount of discriminative information available, and it is likely that more features (i.e. larger) segments will be needed to obtain the same degree of accuracy. Our preliminary analyses using only the limited number of DNAm sites profiled on the EPIC array found that most classifiers offered only a moderate improvement over randomly allocating cell-type labels. This was regardless of whether the classifier was trained to predict five cell-types simultaneously (mean accuracy = 0.34 SD = 0.10) or a series of binary classifiers for each cell-type (mean accuracy = 0.62 – 0.68, **Supplementary Figure 13**). Hence, from here forwards we will only report results from classifiers trained with the WGBS derived DNAm profiles. When predicting five blood cell-types simultaneously, the mean accuracy was 0.75 (SD = 0.15) representing a reasonable improvement over random assignment, with 3,025,646 models (20%) associated with high levels of accuracy (> 0.9). Simplifying the problem to a binary classification problem, there was a mean accuracy of 0.97 (SD = 0.05) for lymphocytes vs myeloid cells, slightly lower than the mean accuracy when trained with continuous DNAm levels (0.99). There was variation in the accuracy of the classifiers for each cell-type with mean accuracy statistics of 0.86 – 0.97 (**Figure 4**). Of note, the monocytes and granulocytes classifiers performed almost as well as the lymphocytes classifiers, with the B-cell and natural killer cell classifiers notably worse. This rank ordering was consistent with what was observed with the classifiers trained on continuous DNAm levels and suggests it relates to how unique the reference profiles for these cell-types are relative to other blood cell-types.

**Figure 4.**
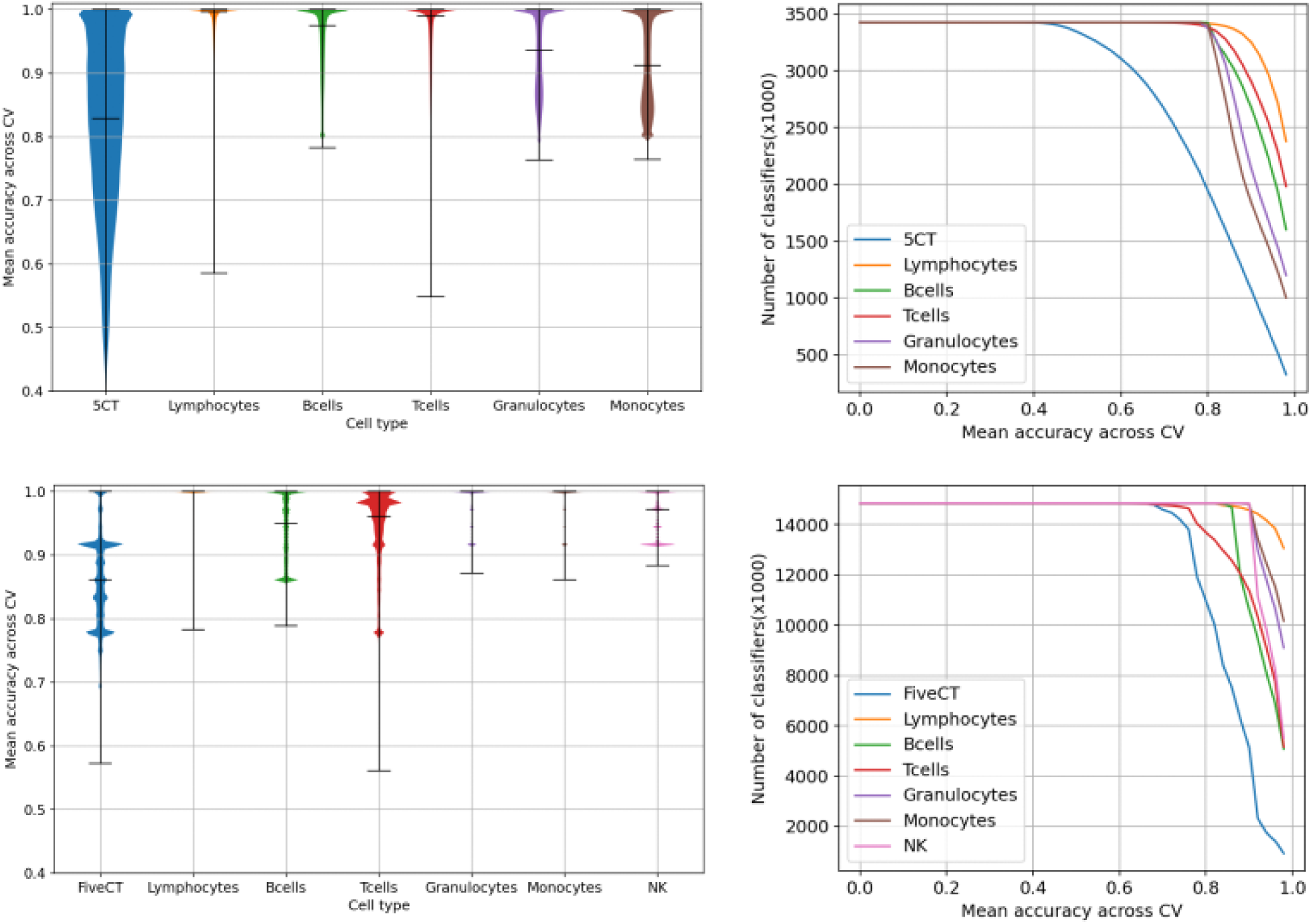
Classifiers can be grouped into a parsimonious set of non-overlapping genomic regions. Plotted are characteristics of non-overlapping set of regions (y-axis) derived from classifiers trained using continuous DNAm levels with accuracy greater than a particular threshold (x-axis). A) total number of regions, B) number of classifiers per region, C) size of each region, D) total size of all regions, E) distance between regions. Each line represnts the mean accuracy score for a different algorithm. Accuracy is defined at the proportion of samples predicted correctly, summarised for each classifier as the mean accuracy score across 15 cross-fold validation iterations.

Profiling how the performance of these classifiers changed as the number of features increased, we observed a very dramatic gain in accuracy for the lymphocyte and monocyte classifiers as the number of sites increased up to 20 before it hit almost perfect performance on average and plateaued (**Supplementary Figure 14**). A similarly dramatic improvement was seen as the size of the classifiers increased to 500bp and peak performance occurred when CpGs were located at an interval of one every 250bp. For the classifier that did not predict as accurately in general, maximal performance did require more information. For example, for the classifiers predicting five cell-types simultaneously, there were valuable improvements up to the inclusion of 100 CpGs before accuracy plateaued. Focusing instead on the most accurate classifiers, again we observed that many of these only required a small number of features. The lymphocyte classifiers (> 0.9) had a mean of 9 sites (just 2 more than that needed for the classifiers built with continuous DNAm levels), and covered a mean of 7.4kb at a mean rate of 1 CpG per 925 bases (**Supplementary Figure 15**). In contrast a mean of 36 sites, spanning 12.4kb were needed to predict five blood cell-types simultaneously. In order to quantify what proportion of the genome contains an accurate classifier we collapsed nested classifiers into regions (**Figure 5A**). For the lymphocyte vs myeloid classifier, 13,634,975 predictive classifiers (accuracy > 0.9) collapsed into 14,053 regions with a mean size of 66.6kb. Altogether, these cover 0.32 of the human genome (**Figure 5D**), demonstrating reasonable potential for the application of this approach for cell-type-specific analysis of DNAm differences.

**Figure 5.**
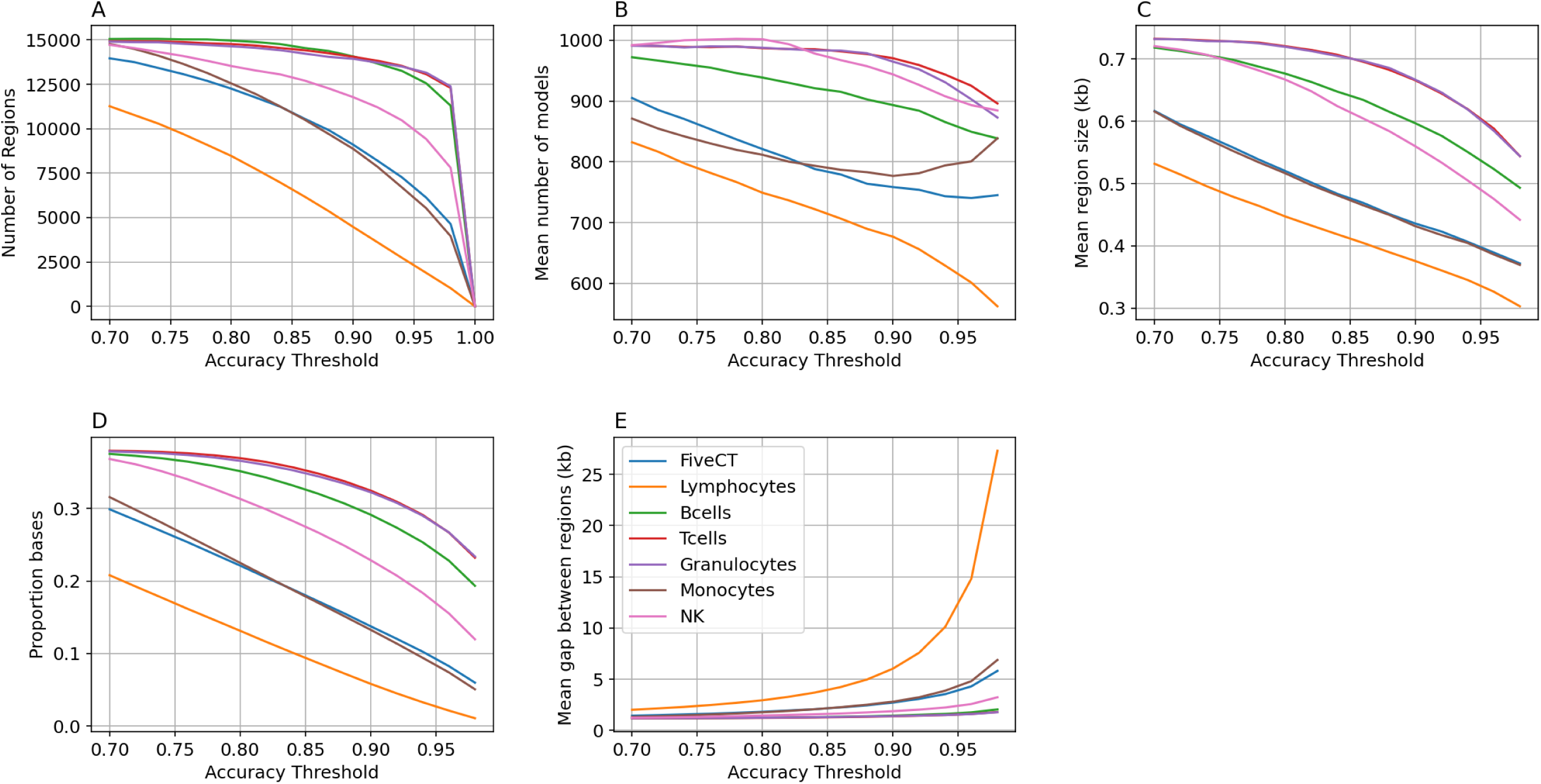
Classifiers can be grouped into a parsimonious set of non-overlapping genomic regions. Plotted are characteristics of non-overlapping set of regions (y-axis) derived from classifiers trained using binary DNAm levels with accuracy greater than a particular threshold (x-axis). A) total number of regions, B) number of classifiers per region, C) size of each region, D) total size of all regions, E) distance between regions. Each line represnts the mean accuracy score for a different algorithm. Accuracy is defined at the proportion of samples predicted correctly, summarised for each classifier as the mean accuracy score across 15 cross-fold validation iterations.

## Discussion

We have demonstrated the potential of using long read sequencing technologies to generate cell-specific DNAm profiles from bulk tissue samples without any additional experiments. By harnessing cell-specific DNAm signatures from a pre-existing WGBS dataset of purified blood cell-types we found that for approximately one third of the genome we could accurately discriminate lymphocytes from myeloid cells. Prediction of more specialised subtypes of blood cell-types was also encouraging but for a smaller proportion of the genome. Our results represent the lower bound of possibility, as the proportion of the genome this approach is viable for will increase as reads get longer and we generate more complete reference DNAm profiles of cell types from long read sequencing.

It is recognised that to understand how the dysregulation of DNAm influences disease aetiology, the analysis of specific cell-types will be required. Ultimately, the goal is to study DNAm at the level of individual cells similar to the way we currently can for gene expression and certain epigenetic modifications. Although methods for single cell bisulfite sequencing(Smallwood et al. 2014; Gu et al. 2021) demonstrate great promise, they are labour intensive, expensive and lack the sensitivity needed for epidemiological studies.

Therefore, isolating populations of cells from bulk tissue is the current gold standard. To perform these investigations, it requires knowing in advance which cell-type(s) to profile and the ability to isolate the cell-type which maybe constrained by the availability of an appropriate antibody or the rarity of that cell type. While for most complex traits, the primary affected tissue may be obvious, the specific cell-type(s) may be unclear, making it challenging to design an appropriate experiment. It is presumed that performing such analysis in the large sample cohorts needed to overcome the multiple testing burden of testing sites genome-wide is experimentally not feasible due to either a lack of existing protocols to isolate the relevant cell-types or the labour and financial demands being prohibitively high. Our computational approach shows that the use of long-read sequencing removes many of these barriers and provides a solution that can be applied to existing datasets to assay potentially multiple different cell-types simultaneously. While we focused on blood cell-types for the purpose of demonstrating the feasibility of the method, our approach can be applied to any tissue or organism where reference data for the constituent cell-types exist. One of our main observations was that the accuracy of cell-type classifiers is not uniform across the genome. This means that each read in a long-read dataset will need to be objectively assessed based on its genomic location, to decide if it contains a cell-type specific signature to confidently classify which cell-type it came from. We suggest that a stringent accuracy threshold is applied as errors in the classification will not only introduce noise into downstream analyses, negatively affecting the statistical power to detect differences, but additionally might introduce false positive associations.

Although application to reads from across the entire human genome is the primary objective, there are situations where this is not necessary. As highlighted earlier one of the challenges for sequencing based DNAm studies is the reliance of read depth or sequencing coverage to quantify accurately the level of DNAm at each genomic position. One way to increase read depth and consequently increase the power of statistical comparisons is to target specific regions of the genome. A number of experimental approaches have been demonstrated to do this successfully in tandem with long read sequencing. Cross-referencing our results prior to the selection of genomic regions to enrich signal from could be a powerful strategy for performing a cell-specific EWAS in the first instance. While our method has been developed for the purpose of disaggregating reads from a bulk tissue, we believe there would also be value in running our method on purified samples, to initially confirm isolation purity, but additionally to identify and remove reads from cells that have been captured erroneously.

While our results suggest that for a large proportion of the genome the classification of long reads by cell-type is not yet possible, as technologies improve and read lengths increase the gaps between the cell-specific regions will be bridged, meaning this is a tractable problem. In the interim, our results provide some highly valuable guidance as to how we can design and train cell-type classifiers to optimise their performance and increase the proportion of the genome for which we can derive cell-specific profiles. One simple way to improve accuracy is to limit the number of cell-types a classifier is trained to predict. While predicting multiple cell-types simultaneously is not an implausible future application, this comes at a cost in terms of the number of features and consequently the length of the sequencing read. It is likely that when predicting more than two cell-types, classifiers need to be bigger because they need to include multiple cell-type signatures across a longer genomic segment. We posit that the more cell-types you attempt to classify, the longer the sequencing read needs to be. Cell-types can be defined at different hierarchical levels with increasing specificity of both function and epigenetic profile. Previous DNAm data has shown that cells from the same linage share a larger proportion of the epigenome and the later they diverge, the more similar they are. Therefore, it becomes an increasingly harder problem to classify more specialised cell-types as there is a decreasing amount of the genome available for discrimination. Carefully considering the level of granularity you truly need or that the data can reliably provide will be an important strategic decision when applying this approach.

A second strategy to maximise the genomic coverage of the predictive classifiers is to profile the reference cell-types using a whole genome sequencing method. Comparisons of sequenced based and microarray-based methods for profiling DNAm have simplified the trade off as increased density of CpGs from sequencing data vs increased precision in quantification of microarray data. Our results suggest that in this application, the increased coverage of DNAm sites captured by the sequencing data outweighs the accuracy of the microarray and means that a smaller genomic segment is needed for classification. It is arguable that WGBS is not truly genome-wide as it affected by the known biases of short read sequencing related to repetitive or CG dense regions. Therefore, we would recommend generating DNAm reference maps with long read sequencing technologies to ensure there is maximum information available to train classifiers.

While our results reveal exciting potential in this area, they should be considered in light of a number of limitations. Firstly, we only used training data from two studies, and it is likely that with the generation of additional cell-type reference the proportion of the genome we can resolve will increase. Secondly, the demographics of the training data (young adult and largely European) might limit the performance in whole blood datasets of other populations. In general we do not think that this is likely given that cell-type differences are the primary driver of variation in DNAm (Hannon et al. 2021; Hannon et al. 2023; Loyfer et al. 2023).

Third, although our method can be easily translated to other tissues and organisms, the specific genomic regions and the proportion of the genome where it works will likely vary. Bespoke classifiers will need to be trained and tested for each new tissue. Fourth, it is dependent on the accuracy of the DNAm calling algorithm from the long read data. This is still an active area of development with multiple algorithms currently available(Liu et al. 2021; Ahsan et al. 2024). Fifth, although the classifiers are capable of identifying multiple different cell-types, the amount of sequencing data you get back will be limited by their abundance.

For example, when classifying reads generated from whole blood derived DNA, while there may be an equal number of accurate classifiers for granulocytes and T-cells, but if granulocytes represent 70% of cells and T-cells 10% of cells, we would anticipate that these percentages would broadly be mirrored in the number of reads classified as each type. This has consequences on the read depth and thus sensitivity in DNAm profiling of each constituent cell-types. For rare cell types very deep sequencing will be required to ensure sufficient sensitivity to detect differences between groups. However, while we have designed an algorithm that does not require knowledge of the cellular composition of the sample to deconvolute the cell-specific profiles, if this information was available and accurate it could potentially be harnessed into the algorithm to improve the accuracy of the classification.

Sixth, we only considered one modification, DNAm, and only then in the context of CpG dinucleotides. For some cell-types, such as neurons, the inclusion of hydroxymethylation will likely increase accuracy of the classification.

In conclusion, we have demonstrated a proof of concept for our computational approach to generate cell-specific DNAm profiles from bulk tissue without the need for additional cell sorting experiments. This advance could unlock the fields understanding of the role of epigenetics in the aetiology of disease by providing the cellular context for associations in population-scale datasets.

## Methods

### Dataset description – microarray derived genome-wide DNAm profiles for purified blood cell-types

The microarray data used in this study are a subset of those described in detail in (Hannon et al. 2021) and are publicly available via GEO accession number GSE103541. Briefly, whole blood samples from 30 individuals from the Environmental Risk (E-Risk) Longitudinal Twin Study(Moffitt and Team 2002; Oliver and Plomin 2007) were collected when the participants were 19 years old. These were subsequently processed using fluorescence-activated cell sorting (FACS) to purify five constituent cell-types (monocytes, granulocytes, CD4+ T cells, CD8+ T cells, and B cells). DNAm was quantified for each sample using the Illumina Infinium HumanMethylationEPIC BeadChip (Illumina Inc, CA, USA) run on an Illumina iScan System (Illumina, CA, USA) using the manufacturers’ standard protocol.

Samples from the same individual were processed together across all experimental stages to negate any methodological batch effects. Following a rigorous quality control process, data was quantile normalised using the *dasen* function from the wateRmelon package(Pidsley et al. 2013). The final dataset for this study included 141 samples and 783,501 sites across the 22 autosomes (**Supplementary Table 1**).

### Dataset description – whole genome bisulphite sequencing derived genome-wide DNAm profiles for purified blood cell-types

These data are a subset of those described in detail in (Loyfer et al. 2023) and are publicly available via GEO accession number GSE186458. Specifically, we used the 36 samples where DNAm was profiled in a blood cell-type. Data was downloaded as beta files (one per sample) and processed within python using custom functions that called on functions in the pandas(McKinney and et al. 2010), statsmodels(Seabold 2010) and numpy(Harris et al. 2020) modules and adapted from the tutorial in wgbs_tools. For each sample we only retained DNAm levels for sites sequenced with a read depth > 10, then filtered to only retain autosomal sites that had DNAm levels for all included samples. We performed an ANOVA for each remaining DNAm site to quantify cell-type differences for across the five major cell groups (granulocytes, monocytes, natural killer cells, B-cells, and T-cells), recording the p-value for filtering later on. The final dataset for this study included 36 samples and 20,368,079 sites across the 22 autosomes (**Supplementary Table 1**).

### Sliding window to select features for classification

Sequencing reads are largely stochastic in nature and can originate from any part of the genome. Long reads can vary not only in the genomic location of the read but also in the length of the read. Therefore, each long read potentially contains a unique combination of DNAm sites and requires a bespoke classifier. To simulate the complete set of possible long reads, and therefore the complete set of epigenetic models consisting of different sets of DNAm sites (i.e. features) required to classify these reads, we used a sliding window methodology. To constrain the number of combinations of DNAm sites, we initially filtered sites based on their evidence for cell-specificity quantified with a statistical test looking for differences in mean DNAm level across sample types. Specifically, we only included sites with an ANOVA p-value < 5x10^-5^. To further constrain the number of different combinations of DNAm sites, we had to train and therefore make the problem computationally feasible and considered a minimum of 5 CpG sites per model and DNAm sites to span a maximum distance of 20,000bp. For each chromosome, sites were the ordered based on their genomic position. The first 5 sites were taken as the features for the first model. We exhausted the search by increasing the size of window to include the next CpG along (a total of 6 CpGs), so long as the span of these sites was less than our predefined maximum distance of 20,000bp. We continually increased the number of sites by one, until our maximum distance was reached. We then “slid” the window across the genome to start at the second site along, shrank the window back to 5 sites and considered all possible classifiers up to 20,000bp before moving to the third site along. We continued in this way along the full length of the chromosome.

### Cell-type classifiers – continuous training data

The classifiers that were developed to use continuous DNAm data as input were trained and tested with the microarray DNAm data for 5 purified cell-types. In this application, we required multiple samples of each cell-type with genome-wide data to enable us to train and test the classifiers. The classification problem was initially built to train a single classifier that predicted all 5 cell-types simultaneously. Training and testing of the classifiers was done using functions from the scikit-learn python module(Pedregosa 2011). For each classifier (defined as a unique combination of features), we used cross-fold validation to quantify accuracy, where the data was split into 3 folds, 2 folds for training and 1 fold for testing, a process that was repeated 5 times using the function *RepeatedStratifiedKFold.* This meant that each classifier was trained and tested 15 times across different splits of the data.

Accuracy was quantified for each iteration as the proportion of correct predictions and summarised as the mean and standard deviation across all 15 accuracy scores. Each classifier was tested using four different supervised machine-learning algorithms: K-Nearest Neighbours (where K was set to the number of cell-type labels in the training data), Naive Bayes, Random Forest, and SVM.

### Cell-type classifiers – binary training data

The classifiers that were developed to use binary DNAm data as input were trained and tested with synthetic data derived from either the microarray DNAm data or the WGBS DNAm data for purified cell-types. To obtain binary (i.e. 0 or 1) methylation status data from levels of DNAm, for each site and cell-type, we sampled from a Binomial distribution with the probability set to the mean DNAm level for that cell-type. The mean DNAm level is a proportion, and therefore falls between 0 and 1 so has the desired statistical properties of a probability. Additionally, it can be considered as the proportion of cells that are methylated and therefore, represents the probability that any single cell picked at random will be methylated. This approach was used to generate multiple synthetic training samples for each cell-type and multiple synthetic test samples for each cell-type. To mirror the cross fold validation approach used for assessing the performance of the continuous classifiers, each classifier was trained and tested on 15 different sets of synthetic data. Accuracy was assessed for each cell-type by calculated the sensitivity and specificity of the predictions in the test data for each of the 15 iterations. These were then summarised by taking the mean across the 15 iterations.

### Software

Data pre-processing of the microarray data was performed using R as described in the original manuscript associated with those data(Hannon et al. 2021). All other analyses were performed in Python3. All code relating to the analyses reported here can be found on GitHub (https://github.com/ejh243/LongReadDNAmCTClassifier).

## Data Access

The microarray derived genome-wide DNAm profiles for purified blood cell-types used in this study are a subset of those described in detail in (Hannon et al. 2021) and are publicly available via GEO accession number GSE103541. The whole genome bisulphite sequencing derived genome-wide DNAm profiles for purified blood cell-types are a subset of those described in detail in (Loyfer et al. 2023) and are publicly available via GEO accession number GSE186458.

## Acknowledgements

E.H is supported by an Engineering and Physical Sciences Research Council Fellowship EP/V052527/1. E.H. and J.M. are supported by Medical Research Council (MRC) grant MR/W004984/1 (awarded to J.M.). This study was supported by the National Institute for Health and Care Research Exeter Biomedical Research Centre. The views expressed are those of the author(s) and not necessarily those of the NIHR or the Department of Health and Social Care. Data analysis was undertaken using high-performance computing at the University of Exeter supported by a Medical Research Council (MRC) Clinical Infrastructure award MR/M008924/1 to J.M.

